# Daraxonrasib (RMC-6236) is an effective targeted therapy for *RAS*-mutant neuroblastoma

**DOI:** 10.64898/2026.02.19.706849

**Authors:** Ronald D. Hill, Krista M. Dalton, Richard Kurupi, Jamie M. Slaughter, Jane Roberts, Yanli Xing, Victor Kehinde, Kun Zhang, Bin Hu, Vita Kraskauskiene, Madelyn R. Lorenz, Jennifer E. Koblinski, Mikhail G. Dozmorov, Konstantinos V. Floros, Anthony C. Faber

**Affiliations:** VCU Philips Institute, Virginia Commonwealth University School of Dentistry and Massey Comprehensive Cancer Center; Richmond, Virginia 23298; Massey Comprehensive Cancer Center and OVPRI, Virginia Commonwealth University School of Medicine, Richmond, VA 23220; Department of Pathology, Virginia Commonwealth University School of Medicine, Richmond, VA 23220; Department of Biostatistics, Virginia Commonwealth University School of Medicine, Richmond, VA 23220; Department of Pediatrics, Virginia Commonwealth University School of Medicine, Richmond, VA 23220

**Author notes:** denotes co-correspondence **correspondence:** Anthony C. Faber, VCU Philips Institute, School of Dentistry, and Massey Comprehensive Cancer Center, Perkinson Building Room 4134, 1101 East Leigh Street, P.O. Box 980566, Richmond, VA 23298-0566., Konstantinos V. Floros, VCU Philips Institute, School of Dentistry, and Massey Comprehensive Cancer Center, Perkinson Building Room 4134, 1101 East Leigh Street, P.O. Box 980566, Richmond, VA 23298-0566.

## Abstract

Neuroblastoma (NB) is the most common extracranial solid tumor in children. Relapsed or refractory (R/R) high-risk (HR) NB tumors continue to exhibit poor outcomes despite intensive and protractive multimodal therapy. Activating mutations in the RAS- mitogen-activated protein kinase (MAPK) pathway are frequently observed in R/R HRNB. The early promise of ALK inhibitors to treat *ALK*-mutant NB underscores the ability of appropriate targeted therapies to improve outcomes for HRNB patients. While MAPK pathway activation is prominent in HRNB, FDA-approved MEK inhibitors and KRAS G12C inhibitors have failed to demonstrate significant preclinical single-agent activity. Daraxonrasib (RMC-6236), a potent and selective RAS(ON) inhibitor, has demonstrated activity in both preclinical models and early phase clinical trials of *RAS*-mutant adult cancers. A subset of R/R HRNB tumors is noteworthy for containing diverse *RAS-*mutations, providing rationale for RMC-6236 investigation. In this study, we evaluated the therapeutic efficacy and oncogenic signaling modulation of RMC-6236 across NB models harboring RAS pathway activation. RMC-6236 as a single-agent treatment led to a significant decrease in cell viability, suppression of downstream MAPK signaling, upregulation of the MAPK pathway effector protein BIM, and increased cell death in *RAS*-mutant NB models as well as in *NF1*-mutant NB models. *In vivo* studies evidenced that RMC-6236 had on-target activity that significantly reduced tumor growth and extended survival in *RAS*-mutant HRNB mouse models. Furthermore, RMC-6236-induced both BCL-2 and BIM upregulation and enhancement of BIM:BCL-2 complexes in *RAS*-mutant NB. As such, the BCL-2 inhibitor venetoclax further enhanced RMC-6236-mediated killing by disrupting RMC-6236 enhanced BIM:BCL-2 complexes. These findings demonstrate that RMC-6236 is a rationale targeted therapy for *RAS*-mutant NB, a subset of NB that is progressively understood as conferring particularly poor outcomes. RMC-6236 is a clinically relevant drug that can successfully target the MAPK pathway in these cancers. This study supports expanded clinical testing of this novel therapy to this important subset of neuroblastoma.

## Introduction

Neuroblastoma (NB) is a cancer of the sympathetic nervous system and accounts for the most cancer-related deaths in children aged five and below^1^, High mortality persists despite improved therapies, including anti-GD2 antibodies ^2^. NB is divided into three risk classes, of which almost all deaths are attributed to the high-risk subset ^3-5^. Specifically, the five-year survival rate for low- and intermediate-risk patients approaches 95%, while the five-year survival rate for HRNB patients is less than 50% ^6-9^. Overall, NB accounts for ∼15% of all pediatric cancer-related deaths, underlying the critical need for surveying promising new therapies that are being investigated in adult cancers. For instance, ALK inhibitors, first tested in *ALK*-mutant NSCLC, have demonstrated efficacy in *ALK*-mutant NB ^10, 11^. Standard therapeutics for HRNB without *ALK* mutations include platinum-based chemotherapy (induction therapy), high-dose chemotherapy plus autologous stem cell transplant (consolidation therapy), and anti-GD2 antibodies like dinutuximab and naxitamab, often times in combination with granulocyte macrophage and colony stimulating factor (maintenance therapy) ^2, 7, 9, 12 31^I-MIBG therapy^13^. isotretinoin^13^, and DFMO.^14^

Alteration of the MAPK pathway is common in many adult carcinomas, including most pancreatic^15^ and a large subset of lung^16^ and colorectal cancers^17^. In NB, the MAPK pathway has also emerged as an important survival pathway, in particular in the R/R HNRB tumors. For instance, genomic interrogation has revealed 80% of R/R HRNB tumors have genomic alterations in the MAPK/ERK pathway ^18-22^. Activating mutations that often present in R/R HRNB include *ALK, KRAS, NRAS*, or *BRAF* genes, while deletion of the RAS GTPase *NF1* (neurofibromin 1) or low expression through non-genetic means, also activates the MAPK pathway in R/R/ HRNB^20, 23, 24^. Higher phospho-ERK (pERK) levels following chemotherapy treatment, as determined by immunohistochemistry, confers poor outcomes^25^.

Underlying the peril of *RAS-mut*ations in NB, Mossé and colleagues recently uncovered that patients harboring HRNB tumors with *RAS-mut*ations at diagnosis with variable allele frequencies (VAF) ≥5% have a distinctively poor 5-year event-free survival (EFS) of 28.6%, versus 46.8% in the HRNB RAS-WT group^18^. Thus, new therapy for *RAS-*mutant NB, which often can indicate ultra-HRNB ^18, 26^, is an important and immediate need.

As a result of these studies and others, inhibitors targeting components of the MAPK/ERK pathway are actively being investigated to improve outcomes in patients with recurrent NB ^11, 23, 27-31^. However, preclinical studies interrogating this pathway with FDA-approved MEK inhibitors like trametinib and binimetinib have revealed disappointing efficacy, due in part to adaptive resistance and feedback activation of alternative growth/survival pathways, including PI3K/AKT/mTOR and FOS/JUN signaling ^29, 32, 33^. Furthermore, a recent investigation of KRAS G12C inhibitors in *RAS*-mutant NBs showed limited activity, likely due to the pan-*RAS-*mutant nature of NBs^34^ as well as the likely importance of wildtype *NRAS* in these cancers^35^. We recently demonstrated^23^ that in *NF1*-mutant and low NF1-expressing NB, SHP2 inhibitors may overcome these resistant mechanisms and provide a therapeutic window that MEK inhibitors do not provide. An important aspect of this enhanced SHP2 inhibitor-sensitivity is they avoid on-target inhibition of MEK in normal-tissue derived cells induced by MEK inhibitors^23^, and thus may have a relatively favorable therapeutic window compared to MEK inhibitors. This hypothesis has yet to be tested clinically in NB. Germane to this study, however, SHP2 inhibitors are ineffective in the vast majority of *RAS*-mutant cancers, including most *RAS*-mutant NB, as active RAS usually circumvents the ability of SHP2 inhibitor to downregulate MEK/ERK signaling^23, 36, 37^.

Covalent KRAS inhibitors designed to inhibit KRAS G12C mutant specific cancers have been immensely successful and have achieved FDA-approval for both KRAS G12C mutant NSCLCs and colorectal cancers^38, 39^. Similar promise has been demonstrated with newer covalent inhibitors targeting G12D, the predominant *KRAS-mut*ation in pancreatic cancer^40^. Differentiating itself from most other cancers, NB includes *RAS-mut*ations that are diversely represented across RAS species HRAS, NRAS and KRAS^18^.

Revolution Medicines has developed a next-generation pan-RAS(on) inhibitor, Daraxonrasib (RMC-6236) ^41^. This drug acts differently from previous generation RAS inhibitors in that it allosterically binds to activated RAS and prevents dimerization with RAF at the RAS Binding Domain (RBD) region, preventing phosphorylation of RAF isoforms and pERK signaling ^41^. RMC-6236 is non-preferential to any RAS isoform or mutation, recognizing and inhibiting all wild-type and mutated forms of *RAS* (*KRAS, NRAS*, and *HRAS*) at relatively equipotent concentrations^42^. Based on significant preclinical activity across *RAS*-mutant NSCLC (non-small cell lung cancer), colorectal and pancreatic cancers^43^, RMC-6236 is in multiple clinical trials in adult cancers, including a phase 3 trial in metastatic pancreatic ductal carcinoma (NCT06625230).

Based on the profile of RAS-mutations in NB and the particularly poor outcomes they confer, the ability of RMC-6236 to inhibit diverse *RAS-*mutants as well as wildtype *NRAS*, and the progression of RMC-6236 to late clinical trials in adult cancers, we assessed whether RMC-6236 had efficacy in *RAS-*mutant and *NF1-*mutant/low HRNB models.

## Results

### RMC-6236 treatment is selectively effective in RAS-mutant and NF1-mutant NB models in vitro

To begin, we treated a panel of NB cell lines with *RAS* activating mutations and *NF1* loss-of-function mutations with RMC-6236 and performed crystal violet (CV) assays^44^ to measure cell viability after treatment with drug concentrations ranging from 0 nM to 100 nM (Fig. 1A, 1B). Treatment resulted in a dose dependent decrease in cell viability across *RAS*-mutant CHP-212 cells (NRAS Q61K)^45^, NB-EBc1 cells (KRAS G12D)^46^, SK-N-AS cells (NRAS Q61K)^46^, LAN6 cells (KRAS G12C)^23^ (Fig. 1A) and NF1-deleted SK-N-FI cells (homozygous deletion)^47^ (Fig. 1B). In addition, sensitivity was expanded to MHH-NB-11 cells which have wild-type *NF1* but profoundly low RNA (DepMap.org) and undetectable protein expression^47^ (Fig. 1B). Indeed, these assays revealed an effective dosage of RMC-6236 in the low nanomolar range. Of note, SK-N-AS cells also harbor a hemizygous deletion in NF1 ^47^. In contrast, RAS WT/NF1 expressing IMR5 NB cells (Fig. 1C) did not display sensitivity to RMC-6236 up to the highest dose tested (100 nM), as would be expected, nor did a panel of normal tissue-derived cells demonstrate any significant sensitivity to RMC-6236 up to 100 nM (Fig. 1D). Thus, both mutations in *KRAS* and *NRAS*, as well as *NF1-*mutant and *NF1* wildtype NB that have pronounced deficiency in NF1 expression, have low (sub 10 nM) nanomolar sensitivity to RMC-6236 *in vitro*, exhibiting roughly two orders of magnitude greater sensitivity than NB cells or normal tissue-derived cells without *RAS*-mutation or NF1 deficiency. Indeed, the sensitivity of these *RAS-*mutants and NF1-mutant/low NBs are analogous to the sensitivity of *RAS-*mutant PDAC in preclinical studies^43^.

**Figure 1.**
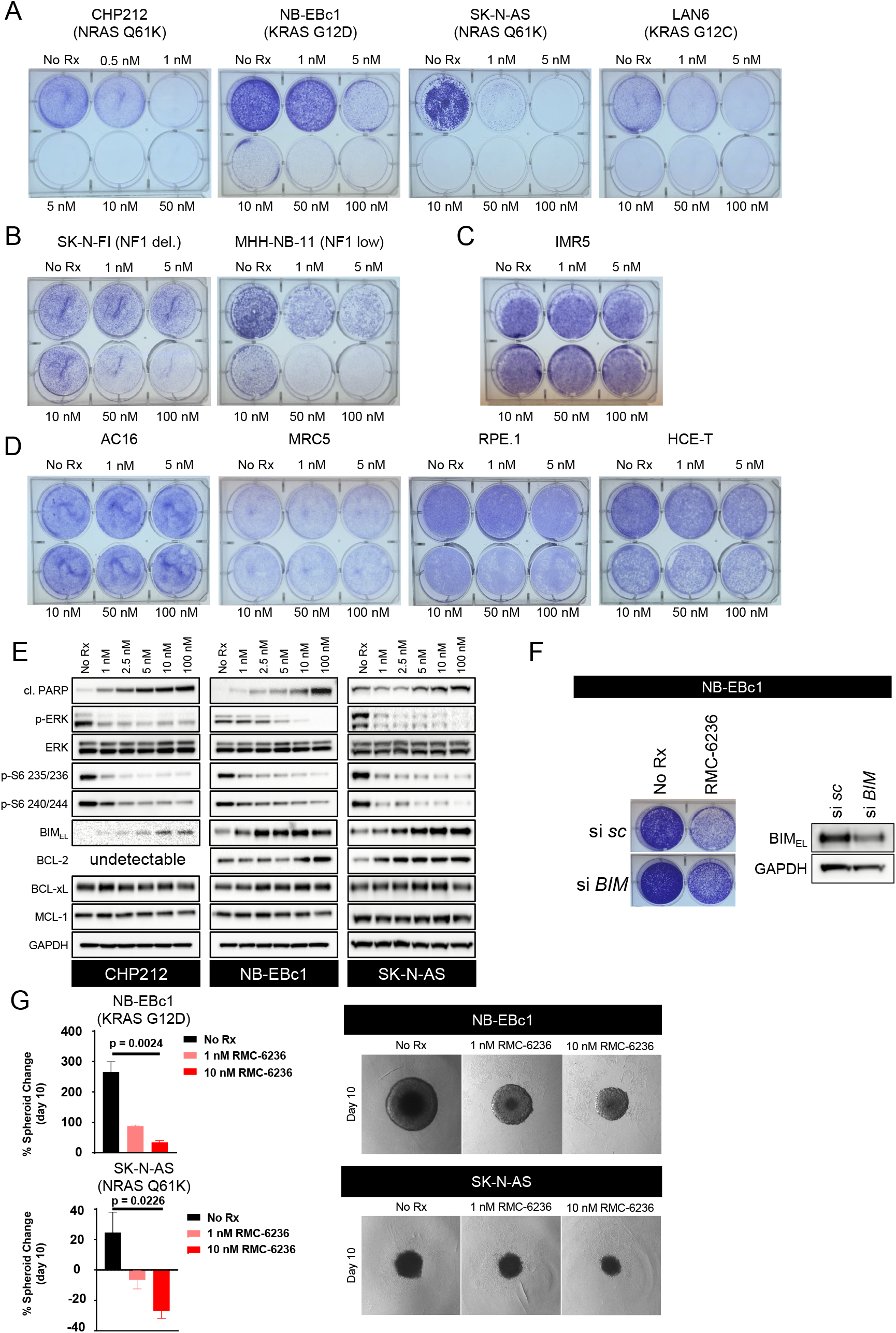
RMC-6236 shows selective efficacy in *RAS*- and *NF1*-mutant neuroblastoma. **(A-D)** Crystal violet staining of NB cell lines harboring *RAS-mut*ations **(A)** and *NF1-mut*ations **(B)**, a *MYCN*-amp cell line (IMR5) with no MAPK-activating mutations **(C)**, and normal tissue-derived cell lines **(D)** treated with increasing doses of RMC-6236 or untreated (No Rx), until No Rx wells reached confluency (typically 5-10 days). **(E)** Western blotting of whole cell lysates previously treated with RMC-6236 for 24 hours and probed for the indicated targets. **(F)** NB-EBc1 cells were transfected with siRNA targeting either a scramble or BIM followed RMC-6236 treatment and crystal violet staining. **(G)** NB-EBc1 and SK-N-AS spheroids were left untreated or treated with either 1 nM RMC-6236, or 10 nM RMC-6236 for 10 days to assess RMC-6236 efficacy in 3D models. Graphs represent average spheroid area change on day 10 compared to day 0 prior to first treatment. Images are of each group on day 10 on the right. (n=3) Error bars are ± SEMs, upper halves shown.

### RMC-6236 suppresses MAPK signaling

To confirm on-target RAS/MAPK inhibition of RMC-6236, we performed western blot analysis across our NB panel. We found RMC-6236 expectedly inhibited the MAPK pathway as evidenced by dose-dependent downregulation of p-ERK (Fig. 1E). In addition, dose-dependent induction of apoptosis, as evidenced by cleaved PARP, was also noted (Fig. 1E). As the mTORC1 pathway is under the control of the MAPK pathway in MAPK-dependent NB, we assessed downstream mTORC1 readouts ribosomal protein S6 (p-S6). We again noted dose-dependent reduction in phospho-Abs (Fig. 1E). BIM is a BCL-2 family protein that is sequestered by anti-apoptotic BCL-2 family members^48^. The MAPK signaling pathway acts to destabilize the BIM protein by targeting it for proteasomal degradation^49^, Consistently, we found BIM was upregulated by RMC-6236 in a dose-dependent manner (Fig. 1E). The anti-apoptotic proteins MCL-1 and BCL-xL did not significantly change with RMC-6236, while BCL-2 levels were increased in the NB-EBc1 and SK-N-AS cells, but was not readily detectable in the CHP-212 cells (Fig. 1E).These data demonstrate RMC-6236 robustly inhibits the MAPK pathway in *RAS*-mutant NBs leading to upregulation of BIM and cell death, despite the upregulation of BCL-2.

To confirm the importance of this MAPK pathway effector protein, we knocked down BIM with siRNA and repeated the CV assay in the presence of RMC-6236 in NB-EBc1 cells. We found genetic reduction of BIM was sufficient to partly rescue the effect of RMC-6236 (Fig. 1F). We recently demonstrated that in NF1-mutant/low NB, mTORC1 signaling is under the control of the MAPK pathway^23^. Indeed, we found in *RAS-*mutant NB, RMC-6236 strongly suppressed the mTORC1 pathway in a dose-dependent manner (Fig. 1E), as evidenced by the reduction of phosph-235/6 and 240/4^50^. Overall, RMC-6236 has on-target MAPK pathway suppression activity, including downstream mTORC1 and BIM.

### RMC-6236 inhibits RAS-mutant NB spheroids

We next determined whether RMC-6236 could block the growth of 3D NB cells. We treated both the NB-EBc1 and SK-N-AS cells with either 1nM or 10 nM RMC-6236 for 10 days. In both models, there was a marked reduction in spheroid area. These data demonstrate anti-NB activity in 3D cultures (Fig. 1G).

### In vivo efficacy of RMC-6236

Given the *in vitro* efficacy of RMC-6236 across these 2D and 3D NB models, we next turned to evaluate RMC-6236 efficacy *in vivo*. Of note, we could not identify any *RAS*-mutant patient-derived xenografts (PDX) models, thus we focused on cell line-derived xenografts (CDXs). We injected these CDX models bilaterally in the flanks of NSG mice, and upon randomization, treated the tumors with vehicle or daily 25 mg/kg^43^ RMC-6236 via oral gavage, a consistent dose with the initial investigations in *RAS*-mutant adult cancers^43^. In the *KRAS* G12C mutant NB-EBc1 model, RMC-6236 attenuated tumor growth significantly over the 4 weeks of treatment compared to the vehicle-treated tumors (Fig. 2A and 2B). Survival analysis demonstrated that by 40d post randomization, all vehicle-treated mice had reached tumor burden limits and were thus euthanized, while all RMC-6236-treated mice were still alive (Fig. 2C), evidencing robust and sustained tumor suppression. These differences were despite drug treatments ending at 4 weeks. Histological analysis of NB-EBc1 excised tumors displayed reduction of Ki67 staining in the RMC-6236 cohort when compared to the vehicle cohort (Fig. 2D). Western blotting of tumor lysates on-target RMC-6236 activity in the tumors, as evidenced by reduced pERK and downstream mTORC1 signaling (pS6) (Fig. 2E), consistent with reduced activation of these pathways in RMC-6236-treated *RAS*-mutant cultured cells (Fig. 1E).

**Figure 2.**
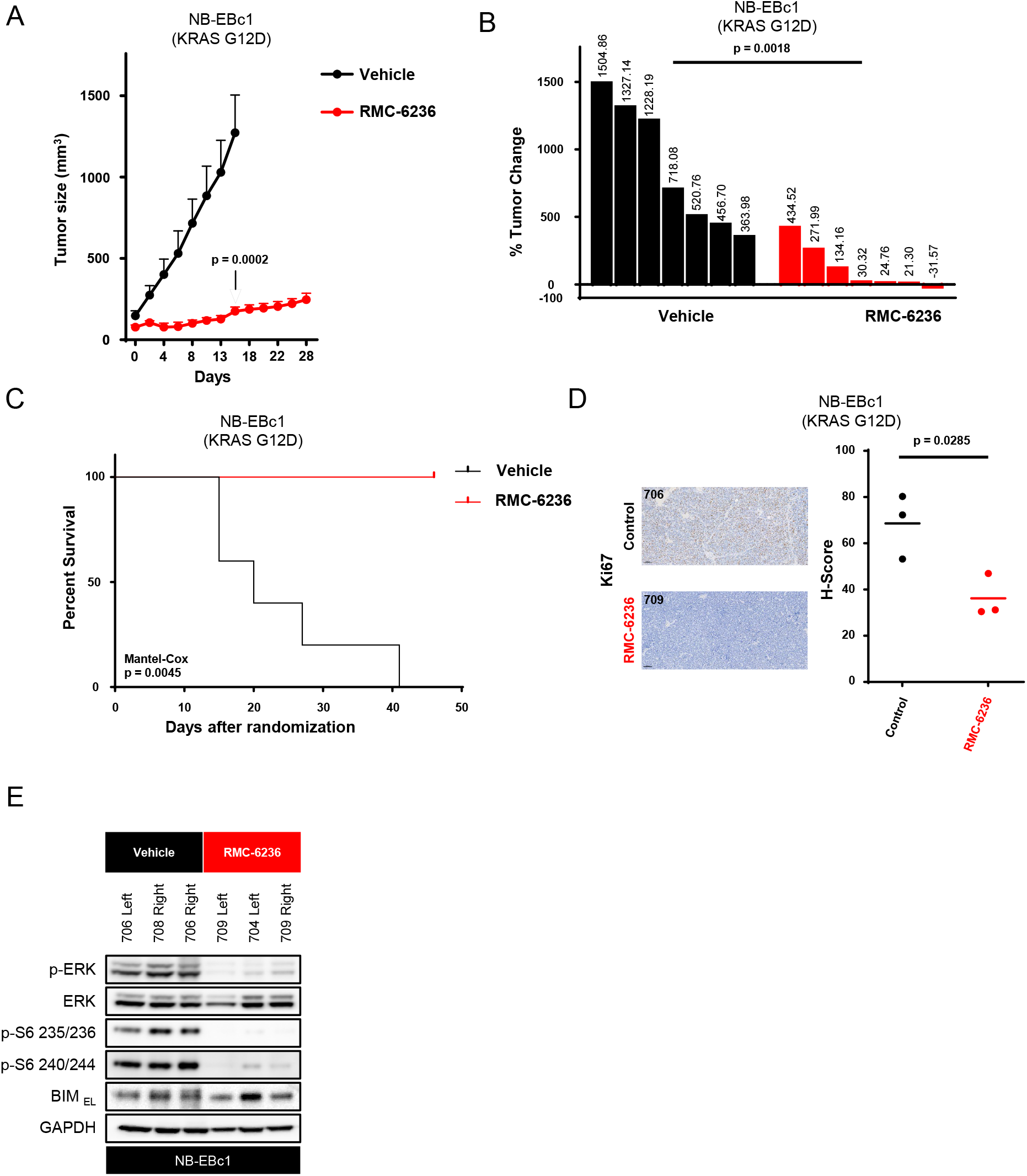
RMC-6236 displays tumor growth attenuation and prolonged survival in *KRAS*-mutant NB. **(A)** NB-EBc1 tumor bearing mice (bilaterally injected) were treated with either vehicle or RMC-6236 (25 mg/kg) via oral gavage for 4 weeks. Tumors were measured via calipers and calculated with the formula v = (l × w × w) (π/6). Unpaired *t* tests were used to calculate significance. **(B)** Waterfall plot represents percent change in tumor volume of each tumor on day 15. Unpaired *t* tests were used to calculate significance. **(C)** Kaplan-Meier curves for overall survival for the duration of the experiment. Mantel-Cox log-rank test was performed to determine significance of survival differences. (n=5 for vehicle and n=4 for RMC-6236 cohorts) **(D)** Representative images of IHC analysis of Ki67 in NB-EBc1 tumors. Scale bars = 100 μm. Histological scores (H-scores) are plotted on the right with bars representing mean value. Unpaired *t* tests were used to calculate significance. **(E)** Western blotting of vehicle and RMC-6236-treated NB-EBc1 tumors probed for the indicated targets. For **A** and **C**, number of mice = 5 for vehicle and 4 for RMC-6236 cohorts. Error bars are ± SEMs, upper halves shown.

In a second CDX, the *NRAS* Q61K mutant SK-N-AS model (Fig. 3A and 3B), RMC-6236 at the same dose and schedule as above, had marked activity, with RMC-6236 shrinking six of the eight tumors (Fig. 3B). Although drug treatment stopped at 28 days, the mice treated with RMC-6236 remained alive several weeks later, while none of the control mice were alive (Fig. 3C). Histochemical scores demonstrated reduced levels in Ki67 the RMC-6236-treated tumors (Fig. 3D). These data together demonstrate robust *in vivo* activity of RMC-6236 in diverse *RAS*-mutant NB models.

**Figure 3.**
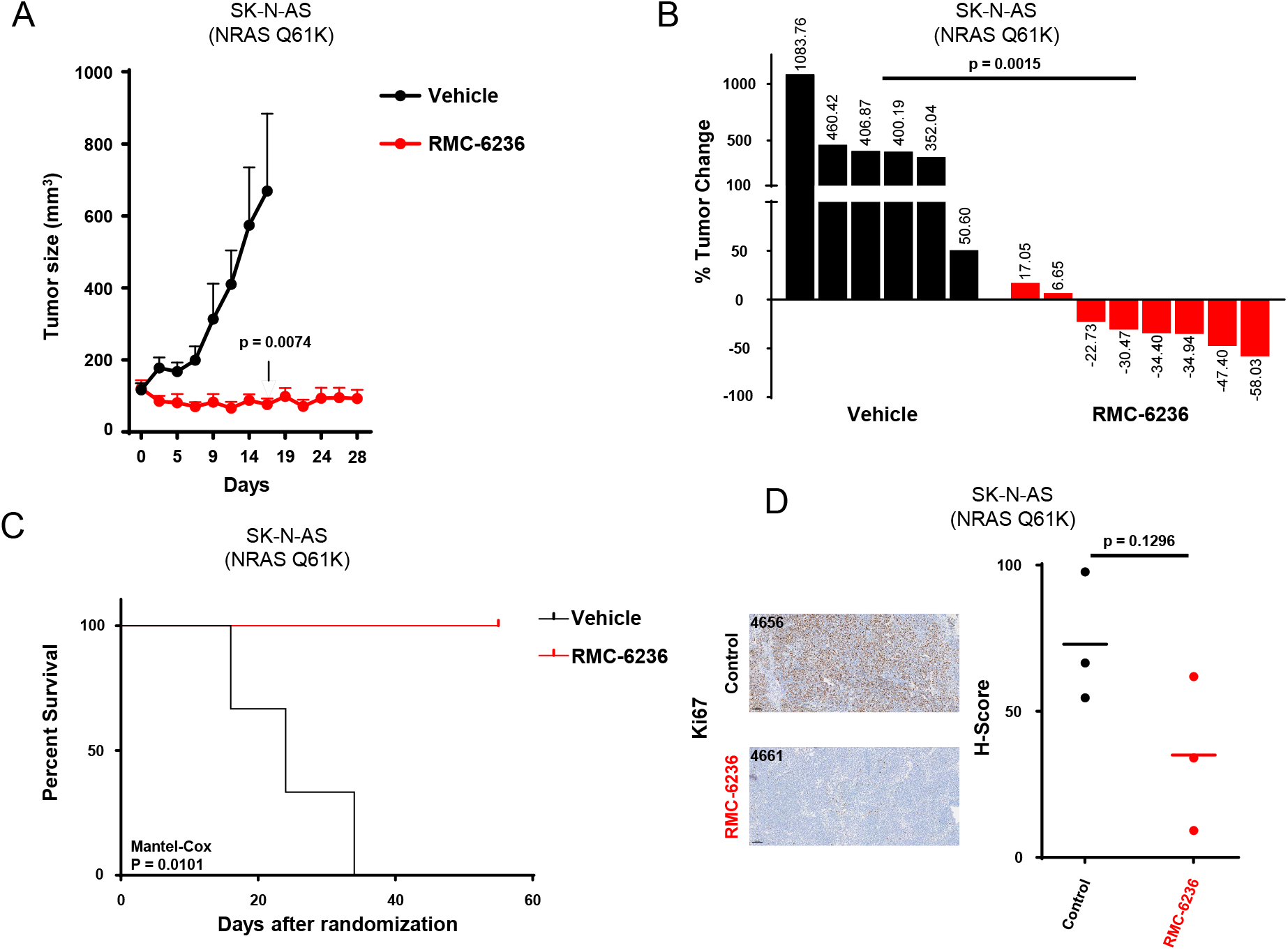
RMC-6236 displays tumor growth attenuation and prolonged survival in *NRAS*-mutant NB. **(A)** SK-N-AS tumor bearing mice (bilaterally injected) were treated with either vehicle or RMC-6236 (25 mg/kg) via oral gavage for 4 weeks. Tumors were measured via calipers and calculated with the formula listed in figure 2A. Unpaired *t* tests were used to calculate significance. **(B)** Waterfall plot represents percent change in tumor volume of each tumor on day 16. Unpaired *t* tests were used to calculate significance. **(C)** Kaplan-Meier curves for overall survival for the duration of the experiment. Mantel-Cox log-rank test was performed to determine significance of survival differences. (n=3 for vehicle and n=5 for RMC-6236 cohorts) **(D)** Representative images of IHC analysis of Ki67 in SK-N-AS tumors. Scale bars = 100 μm. H-scores are plotted on the right with bars representing mean value. Unpaired *t* tests were used to calculate significance. For **A** and **C**, number of mice = 3 for vehicle and 4 for RMC-6236 cohorts. Error bars are ± SEMs, upper halves shown.

### Stabilization of BIM by RMC-6236 primes cells for sensitization by venetoclax

Venetoclax is an FDA-approved BCL-2 inhibitor that is part of standard care for chronic lymphocytic leukemia (CML) and acute myelogenous leukemia (AML). While venetoclax activity is limited across solid tumors, we^51, 52^ and others^53, 54^ have found marked activity across a subset of NB models. Venetoclax killing is largely a BIM-dependent process^55^, and BIM:BCL-2 complexes are a predictor for BCL-2 inhibitor activity^51, 56^. Since RMC-6236 both increased BCL-2 and BIM expression, we asked whether these two drugs would be rationale drug partners. Indeed, we found combination efficacy that exceeded single-agent activity, which was synergistic across several doses of RMC-6236 (Figs. 4A and 4B). In both the NB-EbC-1 and SK-N-AS cells, western blot analysis revealed synergistic killing between the two drugs as evidenced by cleaved PARP and cleaved caspase 3 (CC3) (Fig. 4C). Consistent with the earlier western blot data (Fig. 1E), RMC-6236 increased both BIM and BCL-2 levels, without altering BCL-xL or MCL-1 expression (Fig. 4C). We therefore evaluated BIM:BCL-2 complexes with BIM immunoprecipitation experiments. These experiments demonstrated enhanced BIM:BCL-2 complexes upon RMC-6236 treatment in both models (Fig. 4D), consistent with upregulation of BIM and BCL-2 in whole cell lysates. These complexes were disrupted, as expected, by venetoclax (Fig. 4D). These data together demonstrate that RMC-6236 “primes” *RAS-*mutant and NF1-mutant/low NB for venetoclax-mediated killing through enhanced BIM:BCL-2 complexes, evidencing a mechanistic rationale for combination treatment.

**Figure 4.**
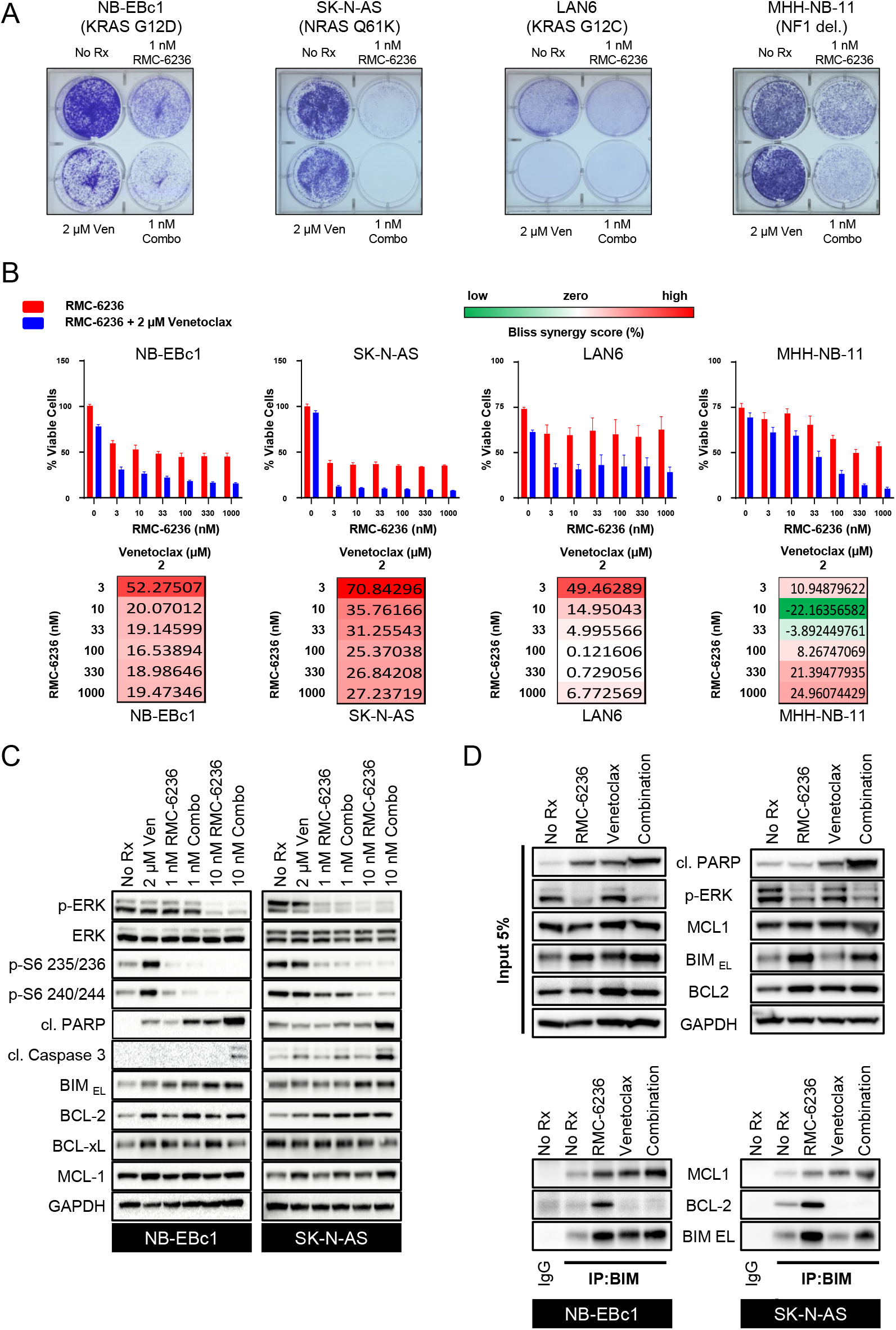
RMC-6236 sensitizes *RAS*-mutant and *NF1*-mutant NB to venetoclax. **(A)** Crystal violet staining of NB cell lines harboring *RAS-mut*ations or *NF1-mut*ations untreated or treated either 2 µM venetoclax or 1 nM RMC-6236, or the combination of venetoclax and RMC-6236 until the No Rx well reached confluency. **(B)** Graphs represent cell viability determined by CellTiter-Glo in the NB cell lines indicated, following 72 hours of treatment with increasing concentrations (0.003–10 μM) of RMC-6236 as shown in red or (0.003–10 μM) of RMC-6236 in combination with 2 μM venetoclax as shown in blue. Bliss synergy scores for concentrations of 0.003–1 μM RMC-6236 with 2 μM venetoclax are plotted below bar graphs. Error bars are ± SEMs. **(C)** Western blotting of NB-EBc1 and SK-N-AS whole cell lysates previously left untreated, or treated with either 2 μM venetoclax, 1nM RMC-6236, 2 μM venetoclax + 1 nM RMC-6236, 10 nM RMC-6236, or 2 μM venetoclax + 10 nM RMC6236 for 24 hours and probed for the indicated targets. **(D)** Immunoprecipitation of whole cell lysates left untreated, or treated with 2 μM venetoclax, 10 nM RMC-6236, or 2 μM venetoclax + 10 nM RMC-6236, for 24 hours. Immunoprecipitation was performed with an anti-BIM antibody. Western blotting was carried out on precipitated and input samples using the antibodies indicated.

### Transient knockdown of BIM prevents apoptosis induced by RMC-6236

SK-N-AS cells were transfected with siRNA targeting either a scramble (sc) or *BIM* sequence followed by treatment of 2µM venetoclax, 10 nM RMC-6236, or a combination of the two drugs. Western blot analysis confirmed BIM silencing and revealed substantial reduction of CC3 in the siBIM combination treated cells (Fig. 5A, *left*), In addition to the control cells (No Rx) growing better, consistent with the characterization of BIM as a tumor suppressor, there was reduced efficacy of RMC-6236, venetoclax, and the combination in BIM-suppressed cells (Fig. 5A, *right*). These data further demonstrate a role of BIM in both RMC-6236 and RMC-6236 + venetoclax toxicity.

**Figure 5.**
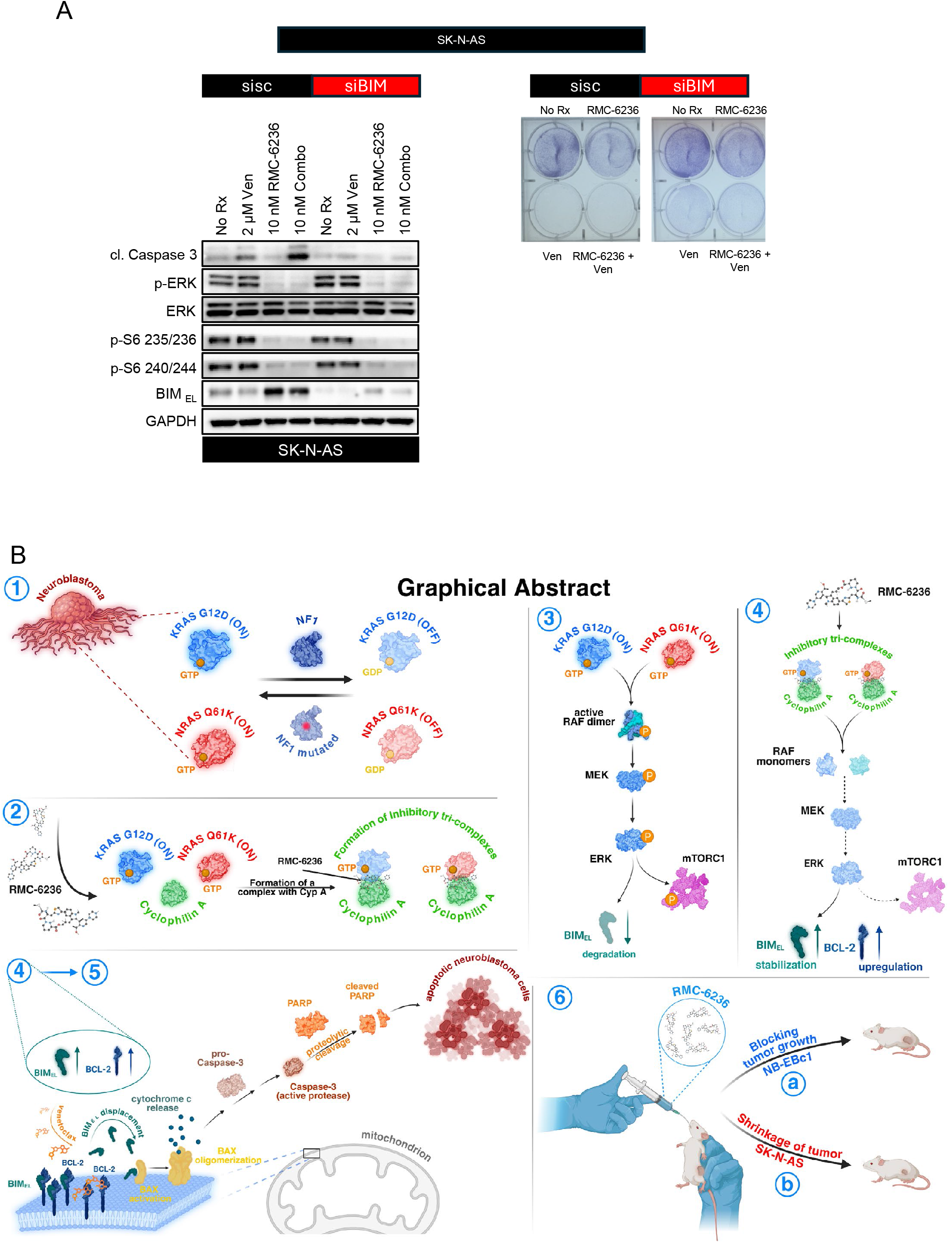
Transient knockdown of BIM rescues NB from RMC-6236 induced apoptosis. **(A)** Western blotting (Left) of SK-N-AS whole cell lysates transfected with either siRNA targeting sisc or siBIM followed by being left untreated or treatment of either 2 µM venetoclax, 10 nM RMC-6236, or 10 nM combination and probed for the indicated targets. Crystal violet staining (Right) of SK-N-AS cell plated after the same transfect and treated with the same drug conditions as western blotting until ‘No Rx’ reached confluency. **(B)** Graphical abstract of the study (1) In neuroblastoma the transition between the GTP-bound RAS (ON) and the GDP-bound RAS (OFF) state can be achieved by the presence of the NF1 WT and the NF1 mutant. (2) After addition of daraxonrasib the cyclophilin A:RMC-6236:NRAS/KRAS tri-complex forms. (3) GTP-bound, active KRAS G12D (NB-EBc1) or NRAS Q61K (SK-N-AS) promotes the MEK/ERK pathway, leading to the degradation of the pro-apoptotic, BH3 only protein, BIM. (4) Treatment with the RAS (ON) inhibitor, RMC-6236, blocks the oncogenic signaling, restores the levels of BIM and upregulates BCL-2. (5) After its stabilization (in 4) BIM is sequestered by anti-apoptotic proteins like BCL-2, on the outer mitochondrial membrane. The addition of a selective BCL-2 inhibitor (venetoclax) displaces BIM, leads to cleavage of PARP and apoptosis. (6) Administration of RMC-6236 in two CDX mouse models (NB-EBc1 and SK-N-AS) results in reduced tumor proiliferation (NB-EBc1) (a) and in tumor shrinkage (SK-N-AS) (b), corroborating our in vitro data (created in BioRender. Floros, K. (2025)).

## Discussion

In this study, we demonstrated that targeting the MAPK pathway with RMC-6236 in *RAS* and NF1-mutant/low NB models HRNB yields promising therapeutic activity. Our findings provide a rationale for clinical investigation of RMC-6236 in these tumors.

RAS pathway mutations, including both *NF1* and *RAS-*mutants, were found to be more prevalent in primary HRNB tumors than primary non-HRNB tumors and confer poor prognosis across all NB tumors regardless of their initial categorization^26^. Additionally, these authors found RAS pathway mutations are enriched in refractory tumors compared to primary tumors^26^. Moreover, HRNB tumors with *RAS-mut*ations at diagnosis with VAF ≥5% carry a particularly poor 5-year EFS of 28.6%^18^.

Despite the clear rationale for MAPK inhibition in HRNB, traditional MEK and ERK inhibitors have shown limited efficacy. Prior work has demonstrated that feedback activation of other growth pathways^57, 58^ contributes to the lack of activity of MEK inhibitor in *ALK*-mutant NB and *RAS*-mutant NB. In addition, we recently demonstrated a lack of therapeutic window of MEK inhibitor between NF1 low NB cells and a diverse panel of normal tissue-derived cells, trametinib was toxic to these models at similar concentrations^23^. Overall, in more general terms, no MEK inhibitor is FDA-approved for any *RAS-*mutant cancers, which likely is a reflection of the lack of therapeutic window that MEK inhibitors provide between these cancers and other tissues, a therapeutic window that is, contrastingly, provided by RAS-specific inhibitors that are FDA-approved across several adult *RAS*-mutant cancers. Recently, cell culture experiments in pediatric cancers including NB exploring KRAS G12C inhibitors indicated a lack of significant activity^34^. This is consistent with the diverse mutations found along the MAPK pathway, and specifically across *RAS* species, in HRNB. In addition, SHP2 inhibitors are ineffective in most *RAS*-mutant cancers, including NB, with the exception of some cancers driven by *KRAS*-specific alleles^23, 59, 60^. Thus, despite the clear rationale to target the MAPK pathway in *RAS*-mutant NB, MEK inhibitors, KRAS isoform inhibitors and SHP2 inhibitors have failed to induce compelling data across *RAS*-mutant NB preclinical models.

RMC-6236 overcomes these limitations and demonstrates consistent preclinical activity across *NRAS, KRAS* and *NF1-*mutant/low NBs. While *NF1-*mutants are only noted in 2-3% of HRNBs, we^23^ and others^47^ have found HRNBs have lower expression of NF1, and low NF1 expression is sufficient to confer poor outcomes. NF1 is reduced likely by from a combination of epigenetic and post-translational changes^23, 47^. For instance, in hemizygous *NF1-*mutant NB cells, the second NF1 allele is often epigenetically silenced^47^. Accelerated proteasomal degradation has also been proposed to reduce NF1 in some NB models^35^, Reduced NF1 expression leads to activation of RAS, and in particular NRAS, the predominantly expressed RAS species in NB^35^. In the MH-NB-11 model, which has wild-type NF1 but markedly low expression of NF1 RNA, RMC-6236 demonstrated low nanomolar activity in these cells (Fig. 1B). Thus, NF1 may be low in a significant number of HRNB, through both epigenetic and post-translational mechanisms, leading to the activation of NRAS/MEK/ERK, and consequently expanding the potential number of HRNB patients beyond those with RAS-mutant and *NF1-*mutant NB that may benefit from RMC-6236.

The emerging clinical data on RMC-6236 has been promising. In *KRAS*-mutant PDAC, RMC-6236 has demonstrated early activity and a manageable safety profile. In addition, circulating DNA has evidenced a marked reduction in mutant *RAS* alleles (ASCO 2025). We found in our study, preclinical efficacy of RMC-6236 was similar in *RAS*-mutant NB (Fig. 1B) to that demonstrated in pancreatic cancer^43^.

Preclinically, a significant subset of NBs, in particular those with *MYCN* amplification^51, 52, 61^ and/or high BCL-2 expression^62, 63^, are susceptible to treatment with venetoclax. This is particularly noteworthy as with the exception of small cell lung cancer (SCLC)^55^, venetoclax is largely ineffective in solid tumors^52^. Clinically, venetoclax in combination with cyclophosphamide and topotecan had encouraging activity in a small cohort of R/R NB patients^64^, substantiating the case that it could be a useful NB therapeutic in appropriate contexts and patients.

Our data provides evidence that RMC-6236 enhances downstream BIM expression, as a result of blocking ERK/1/2-mediated phosphorylation of Ser69 on BIM, marking it for proteasomal degradation^48^. RMC-6236 also upregulates BCL-2, probably as a result of the increased BIM that forms a highly stable complex with BCL-2^65, 66^. This increase in BIM:BCL-2 primes cells for venetoclax-mediated death (Fig. 4D). Venetoclax then disassociates this complex to free BIM to induce apoptosis, providing a molecular mechanism underlying the rationale for further RMC-6236 sensitization with venetoclax. Given that venetoclax has demonstrated tolerability in R/R NB^64^, and it is FDA-approved for several combination therapies in hematological malignancies^67, 68^, this combination provides an intriguing and clinically relevant strategy to further sensitize *RAS*-mutant NB to RMC-6236.

In conclusion, our findings support clinical investigation of RMC-6236 in *RAS-*mutant as well as *NF1-* mutant/low NB. Activity and therapeutic window in *RAS*-mutant NBs both differentiate this RAS-on inhibitor from previously investigated MEK and KRAS G12C inhibitors, offering a targeted therapy option for these HRNB tumors with particularly poor outcomes.

## Study Limitations

Limitations of these experiments include the lack of investigation of RMC-6236 in fully immunocompetent mice, and the lack of orthotopic models.

## Materials and Methods

### Cell Lines

Neuroblastoma cell lines MHH-NB-11, IMR5, LAN6, CHP212, and SK-N-AS were from the Center for Molecular Center Therapeutics Laboratory at Massachusetts General Hospital (Charlestown, MA). The RPE.1, NB-EBc1, and SK-N-FI cell lines were generously provided by Dr. Yael Mosse (University of Pennsylvania, Children’s Hospital of Philadelphia). AC16 cells were kindly gifted by Dr. Fadi Salloum (Virginia Commonwealth University). HCE-T cells were purchased from RIKEN BioSource Center (RBC 2280) and MRC5 cells were purchased from ATCC (ATCC CCL-171). MHH-NB-11, IMR5, LAN6, CHP212, and RPE.1 cells were maintained in RPMI-1640 (Fisher Scientific, cat # 22400105) supplemented with 10% fetal bovine serum (FBS) and 1 µg/mL penicillin and streptomycin. SK-N-AS, SK-N-FI, AC16, and HCE-T cells were maintained in Dulbecco’s Modified Eagle’s Medium/Hams F-12 (50:50) with glutamine (VWR, cat # 10-092-CV) supplemented with 10% FBS and 1 µg/mL of penicillin and streptomycin. NB-EBC1 cells were maintained in Iscove’s Modified Dulbecco’s Medium (IMDM) supplemented with 20% FBS, 1 µg/mL penicillin and streptomycin, and Insulin-Transferrin-Selenium (ITS-G) (ThermoFisher Scientific, cat # 41400045). MRC5 cell line was cultured in DMEM with 10% FBS, 1 μg/mL penicillin and streptomycin. Cell lines were verified via STR testing and routinely tested for mycoplasma infection via Lonza MycoAlert (Fisher Scientific, cat # NC9805874). If found to be positive, cell cultures were treated with Plasmocure (InvivoGen, catalog # ant-pc) per manufacturer’s protocol until cultures tested negative.

### Mouse Models

Tumor Xenograft Models NB-EBc1 and SK-N-AS were injected into the right and left flanks of sex-matched NOD/SCID/IL2Rγ (NSG) mice (aged 5-6 weeks at time of implantation) at a concentration of 5×10^6^ cells per flank for NB-EBC1 and 3×10^6^ cells per flank for SK-N-AS cells. NB-EBC1 were injected into sex-matched male NSG mice while SK-N-AS cells were injected into sex-matched female NSG mice. Cells were injected using a 1:1 ratio of cells to Matrigel (Corning, Cat # 356234). When tumors were measured at a tumor volume of 150-200 mm^3^, mice were randomized to receive vehicle or RMC-6236. Mice in the RMC-6236 treatment group (n=4-5 mice with bilateral tumors) were treated with 25 mg/kg of RMC-6236 via oral gavage while the control group was administered the vehicle via the same route of delivery. Drug was administered for 5 days on, then 2 days off, for a total of 4 weeks. When tumors reached the experimental endpoint of ∼2000 mm^3^ total volume, mice were euthanized via CO^2^ inhalation followed by cervical dislocation. Tumors were then extracted and split into two pieces. One piece was fixed in formalin for histology, and one piece was snap frozen in liquid nitrogen for analysis. All experiments and procedures were pre-approved by the VCU Institutional Animal Care and Use Committee (IACUC) under IACUC protocol AD10001048.

### Reagents

Primary antibodies for western blotting were BCL-2 (cat # 4223S), BCL-xL (cat # 2764S), MCL-1 (cat # 94296S), Bim (cat # 2933S), phospho-S6 Ribosomal Protein (S235/236, cat # 4858S), phospho-S6 Ribosomal Protein (S240/244, cat # 2215S), Erk (cat # 4695S), phosphor-Erk (cat # 4370S), cleaved PARP (cat # 9541S), cleaved Caspase 3 (cat # 9664S); all these primary antibodies were purchased from Cell Signaling Technology (Beverly, MA). GAPDH antibody (cat # sc-32233) from Santa Cruz Biotechnology (Dallas, TX) was used as a loading control. Secondary antibodies used were HRP-conjugated anti-mouse IgG from sheep (cat # NA931-1ML) and HRP-conjugated anti-rabbit IgG from donkey (cat # NA934-1ML) purchased from Cytiva Life Sciences (Marlborough, MA). For immunohistochemistry experiments, antibody Ki67 (cat # 9664) was from Santa Cruz Biotechnology. Daraxonrasib (RMC-6236) was from Abmole (cat # M40832) (Houston, TX) and BOC Sciences (cat # 2765081-21-6) (Shirley, NY), venetoclax was from Medchem Express (cat # HY-15531). Crystal violet was from ThermoFisher Scientific (cat # 42583-0250).

### Western blotting

Cells and tumor extracts were prepared and then lysed in NP-40 containing lysis buffer (20 mM Tris, 150 mM NaCl, 1% Nonidet P-40, 1 mM EDTA, 1 mM EGTA, 10% glycerol, and protease and phosphatase inhibitors). For tumor lysates, mechanical dissociation of the tumor was performed by mincing with scissors followed by homogenization with a tissuemiser (ThermoFisher Scientific) prior to addition of lysis buffer. Upon addition of lysis buffer, cells were incubated on ice for 30 minutes, followed by centrifugation at 16,000 x g for 10 min at 4 °C. Equal amounts of the detergent-soluble lysates were resolved using the NuPAGE Novex Midi Gel system on 4–12% Bis–Tris gels (Invitrogen cat # WG1402BOX), transferred to PVDF membranes (MilliporeSigma cat # IPVH00010) in between six pieces of Whatman paper (VWR cat # 21427-524) set in transfer buffer from Bio-Rad with 20% methanol, and following transfer and blocking in 5% nonfat milk in TBST, probed overnight with the antibodies listed above. Blots were then probed with secondary antibodies conjugated with HRP before adding ECL reagent (ThermoFisher Scientific cat # 34096). Representative blots from several experiments are shown in the figures. Chemiluminescence was detected with the Syngene G: Box camera (Synoptics).

### Spheroid Experiments

For 3D spheroid studies to test RMC-6236 efficacy, SK-N-AS and NB-EBc1 cells were collected at 2.5 x 10^5^ cells and raised to a total suspension volume of 4 mL. 1 mL of SpheroTribe (Idylle Labs), containing 5X methylcellulose solution in DMEM stock was added to each tube of cell suspension. Cell-SpheroTribe suspensions were homogenized by trituration followed by immediate plating of 100 µL of suspension per well in a 96-well round bottom plate (cat # 35117) from Corning. Spheroids incubated at 37°C for 48 hours before imaging and adding drug. All steps were performed according to SpheroTibe manufacturer protocol with the exception of cells plated were half of the recommended count. Images were taken on a Keyence BZ-X series fluorescence microscope (BZ-X810) on days 0 and 10. Spheroid area was calculated using ImageJ. Areas for days 0 and 10 were averaged for each concentration and then graphed.

### Immunoprecipitation

Cells were lysed in the same buffer as was used for western blotting. 500 μg of lysates were incubated each with 2000 ng of Anti-BIM antibody, or 2000 ng of rabbit IgG. 25 μL of 1:1 PBS: prewashed Protein A Sepharose CL-4B beads (cat. no. 17–096303; GE Healthcare Life Sciences) was then added to the antibody/lysate mix followed by overnight incubation with rotating motion. Equal amounts of extracts (5% of immunoprecipitated protein) were also prepared. Representative blots from at least three independent experiments are shown in the figures. Chemiluminescence was detected with the Syngene G: Box camera (Synoptics).

### Histology and immunohistochemistry (IHC)

NB tumor tissues from *in vivo* studies were fixed in 10% neutral buffered formalin which was further suspended in Histogel (cat # HG-4000-012) from ThermoFisher Scientific, followed by processing, paraffin embedding, and sectioning. IHC staining was performed by the VCU Tissue and Data Acquisition and Analysis Core with the Lecia Bond RX autostainer using either heat-induced epitope retrieval buffer 2 (cat # RE7119-CE) from Leica Biosystems, with 45-minute antibody incubation for anti-Ki67 (cat # 9027) from Cell Signaling. Primary antibody incubation was followed by secondary antibody incubation and HRP detection with DAB for 10 minutes using the Bond Polymer Refine Detection kit (cat # DS9800) from Leica Biosystems. Stained slides were then imaged on the Vectra Phenoimager HT (Akoya Biosciences). H-scores were determined by multiplication of the percentage of cells with staining intensity ordinal value (3 × % of cells with 3 + intensity) + (2 × % of cells with 2 + intensity) + (1 × % of cells with 1 + intensity) = H-Score.

### CellTiter-Glo Assays

For CellTiter-Glo (CTG) (Promega) experiments, cells were seeded in quadruplicate in 96-well clear bottom plates at a concentration of 2×10^3^ cells per well in 180 μL of growth medium on day 0. On day 1, cells were treated with increasing concentrations of RMC-6236 from 0 to 10 μM or in combination with venetoclax and maintained at 37°C and 5% CO_2_ for 72 hours. Luminescence was measured on a Synergy H1 monochromator-based multi-mode microplate reader (Biotek) according to the manufacturer’s instructions, except that 25 µL CTG reagent was added to each well before reading rather than the manufacturer’s instructions of 50 µL per well. Non-treated control well values were averaged and normalized to 100 percent viability.

### Bliss Synergy Scoring

The Bliss independence model was applied to evaluate various drugs in combination with RMC-6236. Fractional growth inhibition for each single agent was first determined, and the expected inhibition for a given drug combination was calculated as Expected = A + B – ((A × B)/100), where A and B represent the fractional inhibition values of drug 1 and drug 2, respectively, at the corresponding concentrations. Bliss excess was defined as the difference between the observed inhibition of the drug combination and the Bliss expectation ^69, 70^.

### Crystal Violet Assays

Cells were seeded at a concentration of 0.5 x 10^5^ cells per well in a 6-well plate on day 0. On day 1, cells were treated with increasing concentrations of RMC-6236 0 to 100 nM. Cells were maintained until the control well (No Rx) was confluent as determined by microscope (around 5-7 days) with growth medium containing drug being replenished every 3 days. Once the control reached full confluence, cells were fixed with glutaraldehyde solution (ThermoFisher Scientific) followed by staining with 0.1% crystal violet (Sigma-Aldrich). Plates were then imaged for analysis.

### siRNA experiments

For the siRNA experiments, BIM (Qiagen cat #1027418) and scrambled control siRNA (Qiagen cat # 1027418) were used at the concentration of 25 nM and reverse transfected with lipofectamine RNAiMAX reagent (ThermoFisher cat # 13778075). When doing the transfection, 18 μL RNAiMAX was added to 300 μL of OPTI-MEM (ThermoFisher cat # 31985070) in a 1.5 mL Eppendorf tube. At the same time, 25nM siRNA was added to 300 μL of OPTI-MEM in a separate 1.5 mL Eppendorf tube. After 5 min, the tubes were combined and mixed gently. After an incubation time of 20 minutes, the siRNA mixtures were added to 60 mm dishes. During the incubation period, cells were collected and counted to be plated after siRNA mixture incubation. Cells were then plated on top of the siRNA mixture in 60 mm dishes in antibiotic free medium. 24 hours later, transfected cells were re-seeded into either 96-well plates for CTG, 6-well plates for crystal violet assays, or 6-well plates for western blot analysis. At the time of plating, some cells were collected and lysed to analyze siRNA knockdown efficiency.

## Statistical Analysis

All the methods used to calculate the statistical significance (p-value) for each figure were added to the corresponding figure legends.

## Author contributions

**R.D. Hill:** Conceptualization, Investigation, Data Curation, Formal Analysis, Writing–Original Draft, and Editing. **K.M. Dalton:** Data Curation and Formal Analysis. **R. Kurupi:** Data Curation. **J. Slaughter:** Data Curation. **J. Roberts:** Investigation and Data Curation. **Y. Xing:** Data Curation. **V. Kehinde:** Data Curation. **K. Zhang:** Data Curation. **B. Hu:** Investigation and Data curation. **V. Kraskauskiene:** Investigation and Data Curation. **M. R. Lorenz:** Data Curation and Formal Analysis. **J. E. Koblinski:** Investigation and Formal Analysis. **M. G. Dozmorov:** Data Curation and Formal Analysis. **K. V. Floros:** Conceptualization, Investigation, Data Curation, Formal Analysis, Writing-Original Draft, and Editing. **A. C. Faber:** Conceptualization, Formal Analysis, Resources, Supervision, Funding Acquisition, Writing–Original Draft, Project Administration, Writing–Review and Editing.

## Acknowledgements

This work was in part funded by National Cancer Institute award (R01CA215610-06, awarded to A.C.F.), A.C.F. is also supported by a Natalie N. and John R. Congdon endowment and the Meryl and Charles Witmer Charitable Foundation. Services in support of the research project were generated by the Virginia Commonwealth University (VCU) Cancer Mouse Models Shared Resource and the VCU Massey Comprehensive Cancer Center Tissue and Data Acquisition and Analysis Shared Resource, supported, in part, with funding from NIH-NCI Cancer Center Support Grant P30 CA016059.

